# Hierarchical Deviant Processing in Auditory Cortex of Awake Mice

**DOI:** 10.1101/2023.01.18.524413

**Authors:** Dan Luo, Ji Liu, Ryszard Auksztulewicz, Tony Ka Wing Yip, Patrick O. Kanold, Jan W. Schnupp

## Abstract

Detecting patterns, and noticing unexpected pattern changes, in the environment is a vital aspect of sensory processing. Adaptation and prediction error responses are two components of neural processing related to these tasks, and previous studies in the auditory system in rodents show that these two components are partially dissociable in terms of the topography and latency of neural responses to sensory deviants. However, many previous studies have focused on repetitions of single stimuli, such as pure tones, which have limited ecological validity. In this study, we tested whether the auditory cortical activity shows adaptation to repetition of more complex sound patterns (bisyllabic pairs). Specifically, we compared neural responses to violations of sequences based on single stimulus probability only, against responses to more complex violations based on stimulus order. We employed an auditory oddball paradigm and monitored the auditory cortex (ACtx) activity of awake mice (N=8) using wide-field calcium imaging. We found that cortical responses were sensitive both to single stimulus probabilities and to more global stimulus patterns, as mismatch signals were elicited following both substitution deviants and transposition deviants. Notably, A2 area elicited larger mismatch signaling to those deviants than primary ACtx (A1), which suggests a hierarchical gradient of prediction error signaling in the auditory cortex. Such a hierarchical gradient was observed for late but not early peaks of calcium transients to deviants, suggesting that the late part of the deviant response may reflect prediction error signaling in response to more complex sensory pattern violations.

## Introduction

Deviance detection is a fundamental building block of cognition as it allows for learning and optimizing internal models of the world (Friston, 2005; Auksztulewicz and Friston, 2016; Carbajal and Malmierca, 2018). The mammalian auditory system encodes the same stimuli differently, depending on the context in which they are presented, such as the order of the stimulus in a sequence (Perez-Gonzalez et al., 2005; Yaron et al., 2012; Herrmann et al., 2015), or whether the stimulus is expected or unexpected based on the statistical regularities in the auditory inputs (Dehaene et al., 2015). The classical oddball paradigm has been extensively used for studying the neural mechanisms of deviance detection(Ulanovsky et al., 2003; Naatanen et al., 2007). It involves presenting a sequence of identical repeating stimuli (‘standards’), which as a result have a high probability, and oddball stimuli (‘deviants’), which have a low probability (Cowan et al., 1993). The difference between neural responses to the deviant stimulus and standard stimulus is known as a mismatch response (MMR), and accumulating evidence suggests that MMRs can reflect prediction error signals, rather than mere release from adaptation to the standard(Friston, 2005; Garrido et al., 2009; May and Tiitinen, 2010; Auksztulewicz and Friston, 2015; Carbajal and Malmierca, 2018).

Both humans (Wacongne et al., 2011; Chennu et al., 2016) and macaques(Uhrig et al., 2014; Wang et al., 2015) can detect auditory deviants not only based on probabilities of single stimuli (i.e., in sound sequences of the type …AAAAABAA…, where a particular standard stimulus A has a high probability and deviant stimulus B has lower probability), but also based on more complex patterns involving multiple elements, which may be “chunked” or grouped perceptually according to observed transition probabilities (consider phonemes grouped into syllables and syllables grouped into streams of vocalizations as an example). Similarly, in humans, the auditory cortex elicits MMR also based on unexpected repetitions (…ABABABABAAAB…) and unexpected omissions (…ABABABABA_AB…), which suggests that humans chunk the standard pair (AB) and encode the prediction error to chunk violations (Todorovic et al., 2011; Todorovic and de Lange, 2012; Chouiter et al., 2015). In such contexts, whether individual sounds are expected or not may depend either on their “local” or their “global” context (Chao et al., 2018): e.g., in sequences of the type …AAABAAABAAAB…, each B is locally rare (based on the large number of preceding A) but globally standard (based on the longer stimulus history, with repeated AAAB chunks).

Previous studies in humans and non-human primates suggest a hierarchical gradient of cortical regions encoding local and global deviance (Uhrig et al., 2014; Durschmid et al., 2016; Jiang et al., 2022). However, even invasive methods available in humans such as electrocorticography have a relatively low spatial resolution, resulting in broad distinctions between temporal vs. frontal regions subserving local vs. global deviance processing respectively (Durschmid et al., 2016). On the other hand, in rodents, an extensive literature on mismatch responses to single stimuli suggests a hierarchy of regions also within the auditory cortex, such that secondary areas encode deviance detection to a larger extent than primary areas (Parras et al., 2017; Parras et al., 2021). In addition, an earlier study (Chen et al., 2015) using two-photon guided patch-clamp recordings in the primary auditory cortex of anesthetized mice suggested that the early component (0-100 ms after tone onset) of the neural response to a deviant sound may reflect neural adaptation, while the late component (200 – 400 ms after onset) may reflect deviance detection. However, while rodents can detect single deviants (Ulanovsky et al., 2003; Yaron et al., 2012) and change in acoustic features (An et al., 2021; Yang et al., 2021), to the best of our knowledge there is no direct evidence for MMR signaling in complex sound sequences, although rats can perform sequence chunking if extensively trained (Luo et al., 2021) or if the sequences are repetitive enough (Cappotto et al., 2021). It therefore remains elusive whether the primary auditory cortex, or higher order regions, can also encode deviance in more complex stimulus patterns which are more similar to natural sound sequences.

Here, we aim to test whether the neuronal activity in different regions of the auditory cortex is differentially modulated by distinct types of violation (i.e. at the local element vs. chunk/pair level). To this end, we used continuous, repetitive trains of syllable pairs consisting of artificial syllables (‘pe bi’) as high probability standard stimuli. The syllable tokens were synthesized so as to have formant frequencies that were chosen appropriately for the relatively high frequency range of mouse hearing. The grouping of syllables into pairs sets up a context in which deviations from the expected ‘pe bi’ can be thought of as occurring on a local, within-syllable-pair level (substitutions or omissions) or on a more global level involving the whole pair (transpositions). We then manipulated the elements of the pair to obtain three different types of low probability deviants: (1) substitution deviants (‘pe da’ or ’da bi’), (2) transposition deviants (‘bi pe’), and (3) omission deviants (‘pe _’). Note that the silent interval within a syllable pair (0.145 s) was much shorter than the interval between syllable pairs (2.35 s), which encourages perceptual grouping of the syllable stream into disyllabic “chunks”. As a control condition, we also presented streams of the same syllable pairs (‘pe bi’, ‘pe da’, ‘da bi’, ‘bi pe’ and ‘pe _’) but with equal probability (see Fig. 1A). To obtain neural response to different types of deviations in awake mice, we performed wide-field calcium recordings (see Fig. 1B), imaging from multiple auditory cortical areas simultaneously at a relatively high spatial and temporal resolution. We focused on testing whether cortical activity in the mouse auditory cortex is sensitive to sequence violations, and whether the character of this sensitivity differs for within-pair vs more global, whole-pair violations. Further, we investigated whether mismatch responses to different deviant types differ between primary and higher-order regions.

**Figure 1.**
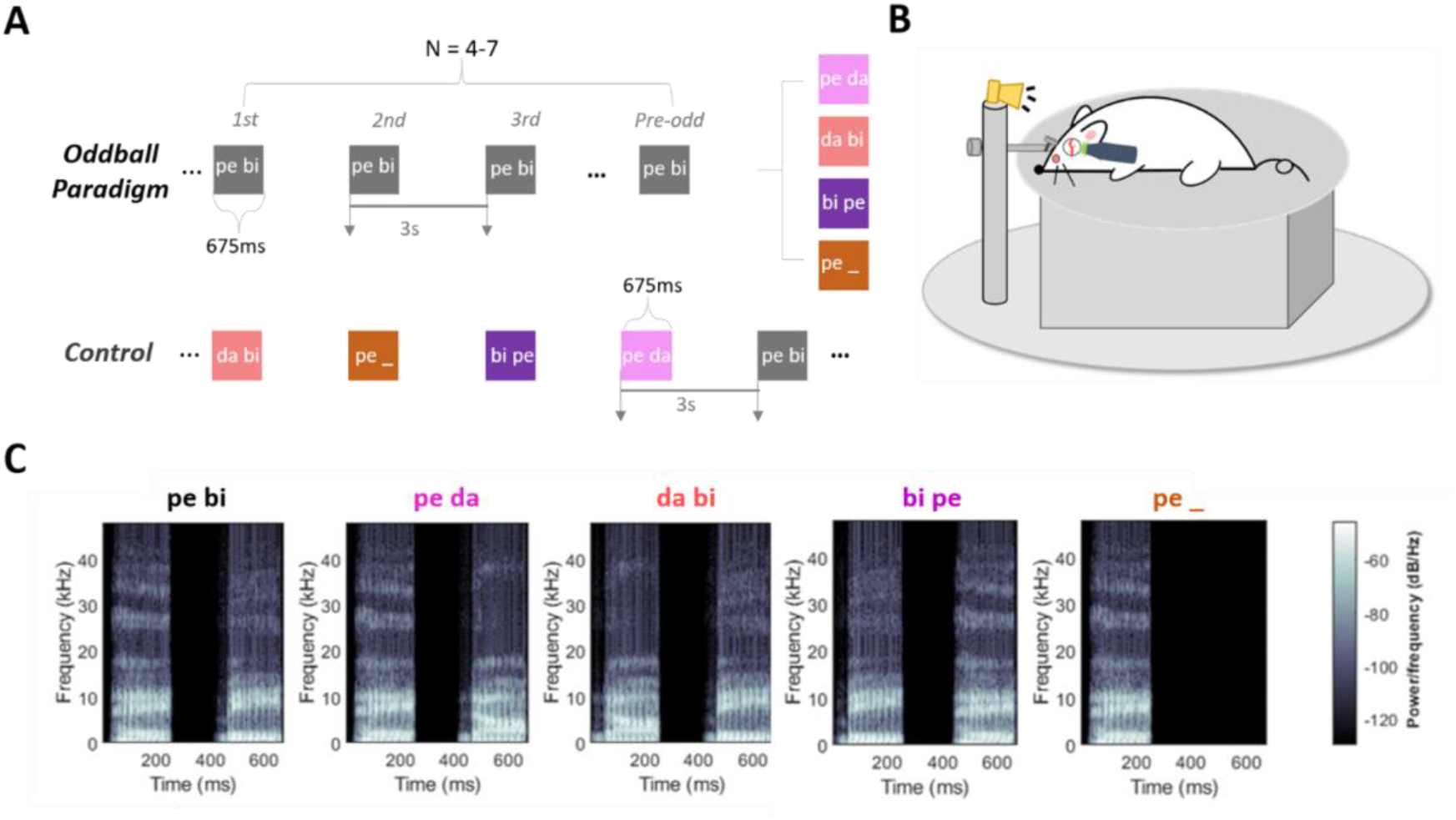
Experimental paradigm, apparatus and auditory stimuli. (A) Schematic of auditory stimuli paradigm. (B) Customized experimental set-up for awake recording. Head-fixed Gcamp6s transgenic mice passively listened to auditory stimuli during wide-field microscopy imaging of the auditory cortex. The mice were placed on a rotatable plastic plate connected with a rotary encoder and the running speed was recorded during the wide-field imaging. (C) Spectrograms of auditory stimuli.

## Methods

All procedures were conducted in accordance with the Guide for the Care and Use of Laboratory Animals and approved by the Institutional Animal Care and Use Committees at the University of Maryland, College Park.

### Animals

Transgenic mice (F1 generation of the CBA/CaJ x Thy1-GCaMP6s mice, mean age = 18.81± 2.58 weeks, 5 male and 3 female) were head-fixed by a customized head holder and placed on a rotatable platform beneath Ultima-IV microscope (Bruker Technologies) during imaging session. The Thy1-GCaMP6s transgenic mouse line is on a C57BL/6 background and expresses the genetically-encoded calcium indicator (GECI) GCaMP6s driven by the Thy1 promoter mostly in excitatory neurons (Feng et al., 2000; Chen et al., 2012; Dana et al., 2014). Since age-related hearing loss is associated with the cdh23 variants in the Thy-GCaMP6s mice on the C57BL/6 background (Kane et al., 2012; Bowen et al., 2020), we used the F1 offspring of the CBA/CaJ mice (JAX000654) and Thy-GCaMP6s (JAX024275) hybrids, which shows normal hearing in middle age (Frisina et al., 2011).

### Surgery for chronic window implantation

We prepared animals as previously (Liu et al., 2019). To prevent brain swelling during craniotomy, 0.1cc dexamethasone (2mg/ml, VetOne, USA) was administered subcutaneously at least two hours before surgery. The animals were initially anesthetized using 4% isoflurane (Fluriso, VetOne, USA) with a calibrated vaporizer (Matrix VIP 3000 Vaporizer, Midmark Corporation, USA). Isoflurane concentration was then lowered to 1.5%–2% during surgery, and the depth of anesthesia was continuously monitored by toe pinch. Body temperature was maintained at 36.0 OC during surgery. Hair remover face cream (Nair, Church and Dwight, USA) was applied on the surgical area (the top and left side of the animal’s head) to remove hair thoroughly. Povidone-iodine and 70% ethanol were applied to clean and sanitize the surgical area. Eye gel was applied to prevent eyes from drying. Then, a longitudinal curved scalp incision was made for skull exposure. Soft tissue under the scalp and periosteum was scraped by a scalpel. A circular cranial window was drilled over the left auditory cortex with a diameter of approximately 3.5mm. The dura mater was gently removed, and gelatin sponges were applied to stop bleeding. To create a chronic optical recording window, a sterile, three layer coverslip was implanted onto the exposed auditory cortex. Surgical silicone adhesive (kwik-sil, World Precision Instruments, USA) and quick adhesive cement (C&B Metabond, USA) were applied to fix the window in situ. Finally, a customized stainless-steel head plate was mounted on the top of the animal’s head using C&B Metabond. This would later allow us to secure the head during imaging sessions. As postoperative prophylaxis, 0.05cc of the antibiotic Cefazolin (1 g/vial, West-Ward Pharmaceuticals, USA), was injected (s.c). After the surgery was completed, the animals were kept individually in the cage and stayed warm under heating lights until they regained consciousness. After recovery, animals are group housed in a reversed 12h-light, 12h-dark light cycle. As further prophylaxis against postoperative infection, the mice received oral antibiotics as a suspension in their drinking water (240mg Sulfamethoxazole and 50mg Trimethoprim in 100 ml water, Aurobindo Pharma, India) for one week following surgery. There were no imaging experiments during recovery (1-2 weeks).

### Experimental design and auditory stimuli

In the FRA mapping blocks, pure tones were generated with custom MATLAB scripts. Each tone lasted 2 s with linear ramps of 5 ms at the beginning and at the end of the tone. Nine tones with equal logarithmic spacing between 4 and 64 kHz were used at three attenuation levels: 0 dB, 20 dB, and 40 dB. Each stimulus was repeated 10 times. Stimuli were presented in a random order.

In the experimental blocks, synthetic acoustic syllable pairs were used to form sequences containing standards and deviants. The syllables were selected from a database of CV syllables recorded by a male speaker (Ives et al., 2005) and were then analyzed and resynthesized by an open-source vocoder, STRAIGHT (Kawahara, 2006), in MATLAB R2018b (MathWorks Inc., Natick, USA). To generate experimental stimuli, we first manipulated and matched the stimulus onset and duration of all syllables (syllable duration = 251 ms), and then shifted the fundamental frequency and formant scalar of each CV syllable upward 1 octave. As can be seen in Figure 1C, the resultant stimuli had a lot of acoustic energy in the 5-40 kHz range, making them well adapted to the relatively high frequency range of mouse hearing (Kelly and Masterton, 1977). In the main experiment (the deviant condition), we used ‘pebi’ as the standard stimulus (365 repeats, 85.88%), and ‘pe da’ (15 repeats, 3.53%), ‘da bi’ (15 repeats, 3.53%), ‘bi pe’(15 repeats, 3.53%), and ’pe_’ (15 repeats, 3.53%) as the deviant stimuli. In the control condition, all stimuli were presented with equal probability and repeated 20 times in a random order. The Stimulus-onset asynchrony (SOA) between pairs was 3 s and the gap between pair elements (offset to onset) was 145ms.

The amplitudes of all auditory stimuli were calibrated to 75dB SPL with a microphone (Bruel & Kjær 4944-A, Denmark). During sound presentation, sound waveform was loaded into RX6 multi-function processor (Tucker-Davis Technologies (TDT)) and attenuated to desired sound levels by PA5 attenuator (TDT). Then the signal was fed into the ED1 speaker driver (TDT), which drove an ES1 electrostatic speaker (TDT). The speaker was placed on the right-hand side of the animal, 10 cm away from the head, at an angle of 45 degrees relative to the midline.

### Widefield imaging

Imaging was performed as previously (Liu et al., 2019). We used 470 nm LED light (M470L3, Thorlabs Inc.) to excite GCaMP6s and acquired the images (camera: PCO edge 4.2, 10Hz, 100ms exposure time, 400 * 400 pixels) with StreamPix 6.5 software (Norpix). Given the single-photon excitation, calcium signals likely predominantly originated from L2/3 neurons (Waters, 2020). We downsampled the original image using the MATLAB built-in function ‘imresize’ from 400 pixels by 400 pixels to 100 pixels by 100 pixels. Next, we performed whitening of the image sequence and image segmentation by using procedures as previously described (Liu et al., 2019), including a constrained autoencoder.

### Trial rejection and data preprocessing

In order to be able to identify periods during which the animals were highly mobile, and during which the quality of the optical recordings would likely suffer from large motion artefacts, we used rotary encoders connected to the plastic plate on which the animals stood to were used for monitoring the movement of the animals. Trial rejection was based on the running velocity of the animal. We detected the bad trials (large movements) by using MATLAB built-in function ‘findpeaks’ and rejected the corresponding trials in the cortical imaging data. For two mice, for which the running speed failed to record, and for those we used an outlier detection to remove putative artifacts, using ‘findpeaks’ to analyze fluorescence values, and setting the outlier threshold ats three times the SD of the median value of the fluorescence. The average percentage of trials rejection was 9.54% across mice (SD=4.54%).

After artifact or outlier rejection, the fluorescence values recorded at each spatial component were normalized in the same manner as described in the previous study (Liu et al., 2019). First, to normalize the raw trace to percentage change, we identified the most frequent fluorescence value F_0_ in a histogram of the whole trace (if there are more than one F values, the minimum F value was selected) and subtracted F_0_ from the whole trace, and then divided the whole trace by F_0_. The F/F_0_ fluorescence trace for each trial was further normalized (z-scored) by the following formula: F =((Ft-mean(F_baseline_))/SD(F_baseline_), where Ft is the raw trace of each trial and F_baseline_ is the baseline period of the raw trace (a 500ms time window before the stimulus onset).

### Spatial components selection

Optical widefield recordings generate time series data for a large pixel array, and the optical values of neighboring pixels are usually very highly correlated. This makes the raw data highly redundant and makes it necessary to include dimensionality reduction steps in the analysis pipeline (Liu et al., 2019). For this purpose, we utilized an autoencoder, following the formulation of the rectified latent variable model (Whiteway and Butts, 2017). Those pixels of the frame with strong temporal correlations across the set of frames were grouped together as individual components. The first 50 components generated by the autoencoder were included for further selection of auditory subdivision for each mouse (Liu et al., 2019). The ROIs of these components covered the entire cranial window of each animal. We then overlaid the FRA map with the spatial map of the autoencoder components for each mouse, together and selected the components that fell within the auditory cortical areas identified by their frequency response fields in the FRA condition separately from each mouse (see Fig. 3D). Components corresponding to blood vessels, as well as those falling outside of the FRA map, were manually excluded. To analyze optical responses across animals but separately for different cortical areas, we collected the selected components for each cortical subdivision across all animals. This yielded 41 components for A1, 26 for AAF and 30 for A2 together for the subsequent analysis (N=8, A1: n=41, AAF: n=26, A2: n=30 components).

### Quantification

To quantify stimulus response amplitudes for each cortical area, we averaged the normalized F values over all spatial components assigned to each of the subdivisions of auditory cortex, and then computed the area under the curve (AUC) of that averaged time series in a 1.5s time window after the stimulus onset. This was done separately for each deviant type and cortical area. To obtain a metric not only of overall calcium responses but also calcium dynamics, we also computed temporal derivatives of the normalized time series Data were also analyzed in the time domain and the temporal derivative was taken for further comparisons by using the matlab function ‘diff’. The first derivative peak values of the average calcium trace was found to match the duration of neuronal firing trains (Carrillo-Reid et al., 2008) and could disentangle the early and late components of the difference curve (Chen et al., 2015). This calculation was done separately for each component and stimulus type in our analysis. Since we identified two distinct temporal derivative peaks to each presented syllable (Fig.5A), we quantified these peaks for the deviant and standard stimuli respectively. We then subtracted the standard peak derivative amplitudes from the deviant peak derivative amplitudes to obtain the “deviant calcium dynamic” (DCD) metric.

## Results

### Parcellation of regions into primary and higher-order auditory cortex

We performed wide-field calcium imaging of the left auditory cortex of 8 adult Thy1-GCaMP6s:CBA F1 mice which express GCaMP6s in excitatory neurons and that have good high frequency hearing into adulthood (Frisina et al., 2011; Liu et al., 2019; Bowen et al., 2020). Before characterizing the cortical responses to syllable stimuli, we first identified the spatial location of different auditory cortical fields by imaging the frequency response areas (FRA) to different frequencies of pure tones (4-64 kHz in half octave steps) at different intensity levels (70, 50 and 30 dB SPL). We found a clear tonotopically organized FRA map for all analyzed subjects (N=8), which showed the expected frequency-dependent gradient in A1, AAF and A2 (Fig. 2B). The fluorescence amplitudes increased most, and larger responsive areas were observed, at 70 dB SPL for most frequencies, while the tonotopic map was much clearer at 30 dB SPL, especially for lower frequencies (Fig. 2A). Based on the relative positions in the mouse brain atlas (Paxinos and Franklin, 2019) coordinates (A1 and AAF are more dorsal than A2, AAF is more anterior than A1, A2 is more ventral than AAF and A1) and the corresponding tonotopic gradients of different areas in the FRA map (Issa et al., 2014; Tsukano et al., 2015; Nieto-Diego and Malmierca, 2016; Liu et al., 2019), we subdivided the imaged auditory cortical areas into A1, AAF, and A2. The tonotopic organization was reliable across all animals (Fig. 2D) and was consistent with previous studies using the GCamp6s transgenic animals (Liu et al., 2019; Romero et al., 2020). To test for functional differences between the three cortical areas (A1, AAF, A2), fluorescence changes were averaged across all components selected for each cortical area. For each subdivision, we selected the components based on the tonotopic map of each mouse from the first fifty components generated by an autoencoder analysis (Liu et al., 2019) and merged the components from all mice for the subsequent analysis.

**Figure 2.**
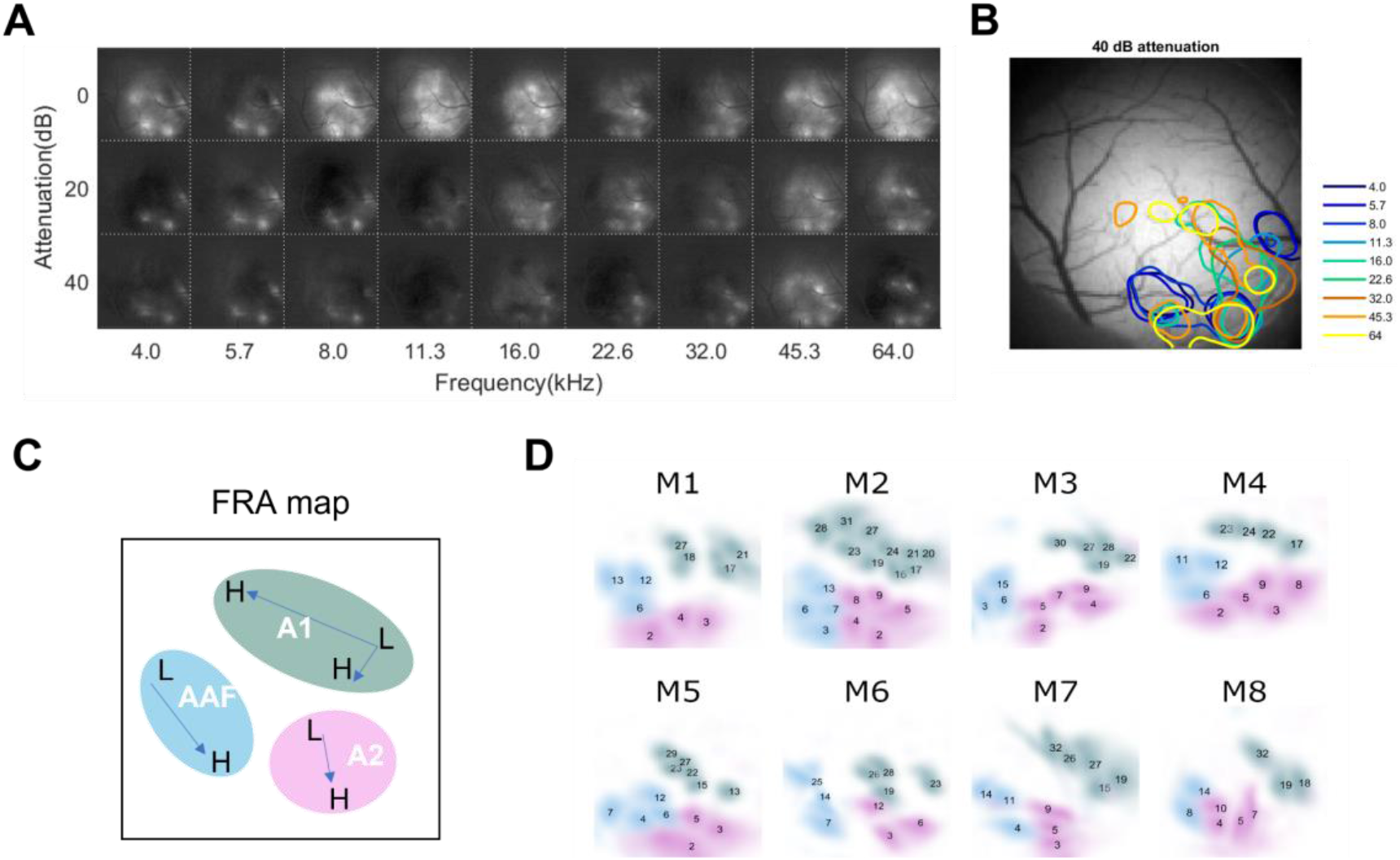
Identification of the distinct auditory fields by wide-field calcium imaging. (A) Example of WF calcium signal to pure tones of different frequencies and sound levels of pure tones in one animal. Each image shows the averaged evoked response within 200-500 ms after tone onset. (B) Example of tonotopy map at 30 dB SPL in one animal. Different frequencies (kHz) of pure tone are color-coded, and each contour line illustrates the response area to the corresponding frequency. (C) Schematic of a tonotopic FRA map. Arrows represent gradients of low to high frequency preference. Distinct auditory fields (A1, AAF, A2) marked in different colors. (D) Tonotopic FRA maps of 8 animals. Color indicates the corresponding auditory fields. Numerical values indicate the selected components based on the FRA map of each mouse.

### Quantification of adaptation and mismatch responses

Having mapped the subdivisions of the auditory cortex, we first quantified neuronal adaptation by analyzing the responses to subsequent standard syllables. We qualitatively observed that the fluorescence of the response to standard stimuli decreased from the first standard to the last repeat (immediately preceding the oddball stimulus; “pre-odd”) in A1, AAF and A2 (Fig. 3A). To quantify the neural adaptation effect in each auditory cortical area we performed linear regression on the area under the curve (AUC) of the fluorescence in response to the first three standard repeats and the last repeat. We identified a significant decrease of the AUC values from the first repeat to the last repeat in all areas (A1: F(2,162) = 10.7, P = 0.001; AAF: F(2,102) = 4.46, P = 0.037; A2: F(2,118) = 4.93, P = 0.028; see Fig. 3B), suggesting that adaptation to repeating standards occurs across all imaged areas.

**Figure 3.**
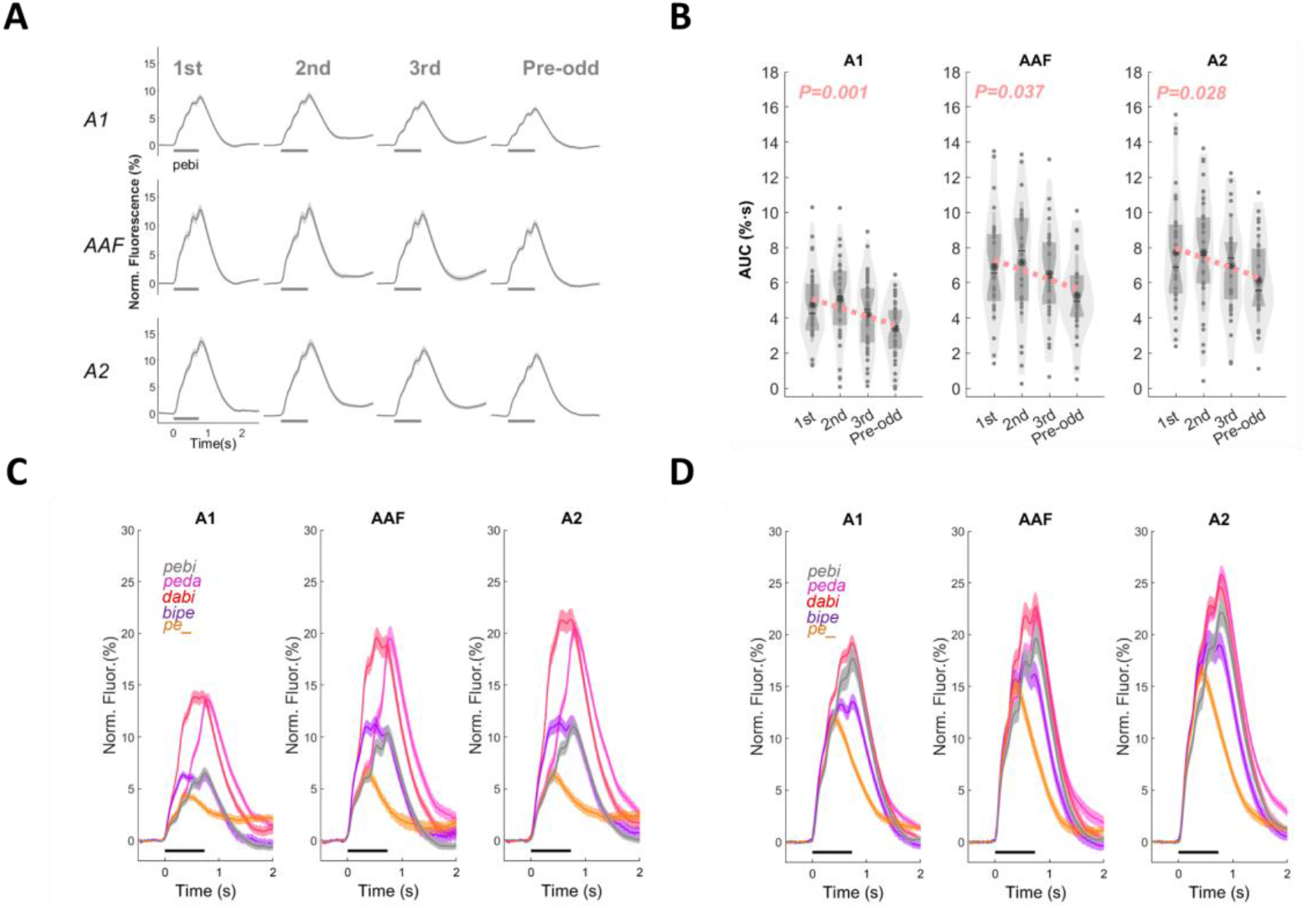
Adaptation and mismatch responses. (A) Neural adaptation to subsequent repetitions of standard stimuli. The calcium fluorescence traces show the averaged response to the standard stimuli (at the first, second, third, and the preceding position before the deviants in each repeating cycle) over all selected components in different auditory fields (A1, AAF, and A2). Dark horizontal bars below the traces indicate when the stimuli were playing. (B) Linear regression quantifying adaptation. Each violin plot shows the distribution of the area under curve values of the fluorescence signals attributed to changes in neural activity in each of the three cortical fields by the autoencoder analysis. Each black dot shows AUC values from one component in that auditory field. Asterisk indicates the mean of the AUC values in that cortical area. Pink dashed lines show linear regression and p-values indicate the significance of the linear regression. (C) Calcium response to deviants and standards in oddball condition. Color coded line presents the calcium fluorescence trace to different types of auditory stimuli averaged over selected components (pooled across animals) in each auditory field. The dark bars indicate the stimulus period. (D) Calcium responses to the same stimulus types in the control condition, plotted like in panel (C).

Then, we proceeded with quantifying deviance effects. As an initial step, since different cortical regions might be differentially sensitive to the syllables which formed the standard (‘pe bi’) and deviant stimuli (e.g., ‘pe da’), we tested whether the syllable pairs differed significantly in how strongly they activated cortex, independently of oddball or frequency effects. To this end, we performed two-way ANOVAs on AUC values of fluorescence. We tested for differences between the two main stimulus types (the standard vs. all deviants pooled together) and auditory areas (A1, AAF, A2), separately for each of the oddball conditions and for the control condition in which all stimuli were presented with equal probability.

In the control condition, there was no effect of stimulus type (*F* _(1,188)_ = 0.27, p=0.605) but there was an effect of cortical area (*F* _(2,188)_ = 13.51, p < 0.001). Post-hoc Bonferroni tests showed that the overall response magnitude in A2 was larger than in A1 (p < 0.001) or AAF (p = 0.013), but there was no difference between A1 and AAF (p = 0.172). Besides, no interaction effect between stimulus type and areas was found in the control condition (F (2,188) = 0.12, p = 0.889). These results indicate that the population of neural activity in auditory cortex is relatively similar to the different stimuli used here when they were matched for presentation probability.

In contrast, in the oddball condition, we found an effect for stimulus type (*F* _(1,188)_ = 61.78, p < 0.001) as well as for auditory area (*F* _(2,188)_ = 41.57, p < 0.001), but no interaction (*F* _(2,188)_ = 0.28, p =0.757). Post-hoc tests revealed a difference between deviants and standards (p < 0.001), showing that the deviants evoked a larger response in auditory cortex to the deviants than the standards, as would be expected given the extensive literature on stimulus specific adaptation (Ulanovsky et al., 2003; Yaron et al., 2012). Furthermore, we found the overall response in A2 and AAF to be stronger than in A1 (Bonferroni test, both p < 0.001), with no difference between AAF and A2 (p = 0.138).

Taken together, we found evidence for a general gradient of response amplitudes to acoustic stimuli, with relatively stronger responses in higher-order areas such as A2. Crucially, these results also demonstrated that deviant responses were stronger than standard responses in the oddball condition, but not in the control condition. While pooling all deviant stimuli together did not reveal an overall difference in mismatch responses across the imaged areas, we reasoned that such differences may arise when comparing responses to different types of deviants individually. Therefore, in subsequent analyses we focused on the different oddball conditions and tested whether different types of deviants modulate the amplitude of mismatch responses, and whether this modulation differs between auditory regions.

### Robust differences in mismatch signaling between the non-primary area (A2) and A1 area

To estimate the mismatch responses evoked by different types of deviant stimuli (Fig. 4A), first we compared the AUC of fluorescence to the substitution (‘pe da’, ‘da bi’) and transposition (‘bi pe’) deviant stimuli against the pre-odd standard stimuli (‘pe bi’) in oddball blocks. The false discovery rate (FDR < 0.05) was applied to correct for multiple comparisons across tests. This analysis showed that, for deviants containing a substitution of the second element of the pair (‘pe da’), the AUC was larger for oddball stimuli compared with the pre-odd standard stimuli. This was true in all auditory fields (paired t-tests; A1: t_41_ = 14.084, p < 0.001; AAF: t_25_ = 9.832, p < 0.001; A2: t_29_ = 13.411, p < 0.001; FDR corrected). Similarly, for deviants containing a substitution of the first pair element (‘da bi’), we found the same pattern of results in all areas (A1: t_41_ =15.138, p < 0.001; AAF: t_25_ = 10.126, p < 0.001; A2: t_29_ =10.342, p < 0.001; FDR corrected). Interestingly, for the transposition condition (‘bi pe’), we found significant differences between the AUC of the deviants and the pre-odd standard in the AAF and A2 areas (AAF: t_25_ =3.031, p = 0.006; A2: t_29_ =3.224, p = 0.003; FDR corrected; compare middle and third columns in Fig 4A), but not in the A1 area (p > 0.05). These results suggest that A1 shows strong mismatch responses to deviance based on the substitution (likely due to adaptation, as mismatch responses are here calculated relative to the pre-odd standard), but not to deviance based on transposition; conversely, higher-order areas A2 and AAF show mismatch responses to both kinds of deviance manipulations.

**Figure 4.**
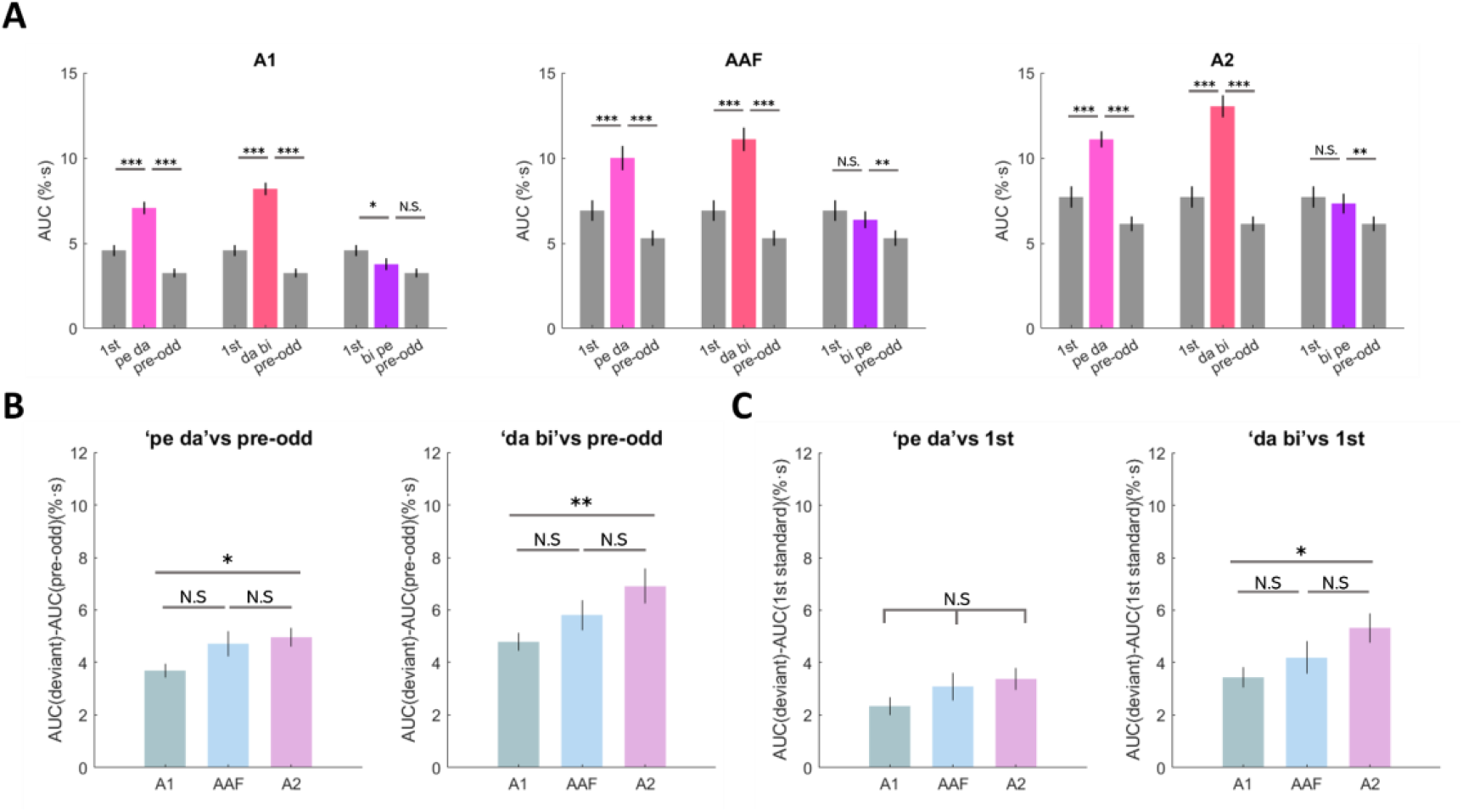
Response differences between deviants and standards across deviant types and auditory fields. (A) Comparison of calcium response to the first standard stimuli, pre-odd standard stimuli and oddball stimuli across different deviant types and distinct auditory fields. (B) Between-area comparison of mismatch response to the deviant (AUC_deviant_) and the responses to the standard preceding the deviant(AUC_pre-odd_) between cortical areas in oddball condition. (C) Comparison of mismatch response to the deviant(AUC_deviant_) and the responses to the first standard following the oddball (AUC_first_) between cortical areas in oddball condition.

In the above analysis, we compared the responses to deviants against responses to an adapted pre-odd standard. Next, we sought to quantify the mismatch response after taking adaptation to the standard into account. To this end, we compared the AUC of fluorescence to the deviant stimuli (‘pe da’, ‘da bi’, ‘bi pe’) against the first standard stimuli (‘pe bi’) in oddball blocks. The results showed that for substitution deviants (‘pe da’ and ‘da bi’), the AUC of response to deviants was larger than that to the first standard stimuli in all auditory fields. This was the case for both substitution deviants: ‘pe da’ (paired t-tests, FDR corrected across tests; A1: t_41_ = 7.32, p < 0.001; AAF: t_25_ = 5.836, p < 0.001; A2: t_29_ =7.990, p < 0.001) and ‘da bi’ (A1: t_41_ =9.518, p < 0.001; AAF: t_25_ = 6.798, p < 0.001; A2: t_29_ =9.646, p < 0.001). On the other hand, for the transposition deviant (‘bi pe’), we found that the AUC of the deviants was smaller than that to the first standard in A1 (t_41_ =-2.633, p = 0.012; FDR corrected), but not in AAF or A2 (both p > 0.05). Please note that the latter result shows a decrease in A1 AUC for transposition deviants relative to the first standard, suggesting that A1 does not signal a mismatch response to transposition deviants (as quantified with AUC), but rather adaptation to the repeated syllables independent of their order.

Having shown that there are mismatch responses to the substitution deviants in all fields, independently of the reference standard (deviants vs. first standards as well as pre-odd standards), as well as mismatch responses to transposition deviants in AAF and A2 when comparing deviants vs. pre-odd standards, we then tested for differences in mismatch responses between fields. Due to the sample size differences between different cortical areas, rank-sum test was used for the comparison over different auditory fields of the mismatch response (deviant against the first standard or against the pre-odd standard) in two substitution deviant conditions respectively (‘pe da’, ‘da bi’). To this end, for each area and deviant type, we subtracted responses to standards from responses to deviants, and entered the difference values into rank-sum tests between cortical areas. The larger deviant response was found in A2, especially to the substitution deviants. We found there was a significant difference between A1 and A2 in mismatch responses to substitution deviants compared with the pre-odd standards (rank-sum test; Z_*peda*_ = -2.520, P_*peda*_ = 0.012; Z_*dabi*_ = -2.858, P_*dabi*_ = 0.004; FDR corrected; see Fig.4B), but not between A1 and AAF (rank-sum test, all P > 0.05, FDR corrected; see Fig.4B), or between AAF and A2 (rank-sum test, all P > 0.05, FDR corrected; see Fig.4B). However, when analyzing mismatch responses relative to the first standard, we only identified a significant difference between A1 and A2 in mismatch responses to deviants based on substituting the first element of the pair (‘da bi’; mismatch response significantly stronger in A2, rank-sum test, Z = -2.427, P = 0.015, FDR corrected; see Fig.4C), but not between other pairs of regions, and not for other types of deviants (all other P > 0.05;see Fig.4C).

### Late calcium responses differentiate between deviant types and auditory areas

Given previous reports that the roles of neural activity at different latencies might be functionally dissociable (Chen et al., 2015), we also wished to perform further analyses at finer time resolution than the AUC based quantification over the entire response. To this end, we calculated the first temporal derivative of the fluorescence traces and subjected them to further analysis (see Fig. 5). The rationale for considering derivatives is that time periods during which the calcium signal increases rapidly (large temporal derivative) should correspond to periods of strong, response related neural excitation. In the temporal derivative time series, we observed four distinct response peaks (i.e., an early and a late response to each syllable in a pair), based on which we specified four distinct time windows for the sub-syllabic responses: the early peak of the first syllable (0-133 ms after the first syllable onset), the late peak of the first syllable (166-300 ms after the first syllable onset), the early peak of the second syllable (0-133 ms after the second syllable onset) and the late peak of the second syllable (166-300 ms after the second syllable onset). Of those four peaks, only two, either the first or the second pair, were considered of particular interest in each case. For the ‘pe da’ substitution, the first syllable ‘pe’ is the same in the standard and the deviant, so only the derivative peaks associated with the second syllable can exhibit deviant responses. Fig.5A indicates that this is indeed the case. In contrast, for the ‘da bi’ and ‘bi pe’ deviants, the auditory pathway may be able to detect the deviation from the standard already during the first syllable, so in those cases we look for evidence of deviant responses in the first pair of peaks. To quantify deviance responses in these respective peaks, we compared the peak amplitudes of the responses to the deviant elements identified above (peaks of red or purple curves in Fig.5A) against the responses to the pre-odd standard elements in the same positions within the curve (black curves in Fig.5A). For example, for the ‘da bi’ deviant (vs. ‘pe bi’ standard) we compared the peaks corresponding to the deviant element ‘da’ (first syllable) against the peaks corresponding to the standard element ‘pe’ (first syllable). The difference in these peak amplitudes we refer to as the “deviant calcium dynamic” (DCD) metric. Given that early and late peaks have been hypothesized to have different functional roles (Chen et al., 2015), we have calculated DCD values separately for the initial peak (early time window; pink shaded area in Fig. 5A) and the later peak (late time window; cyan shaded area in Fig. 5A).

**Figure 5.**
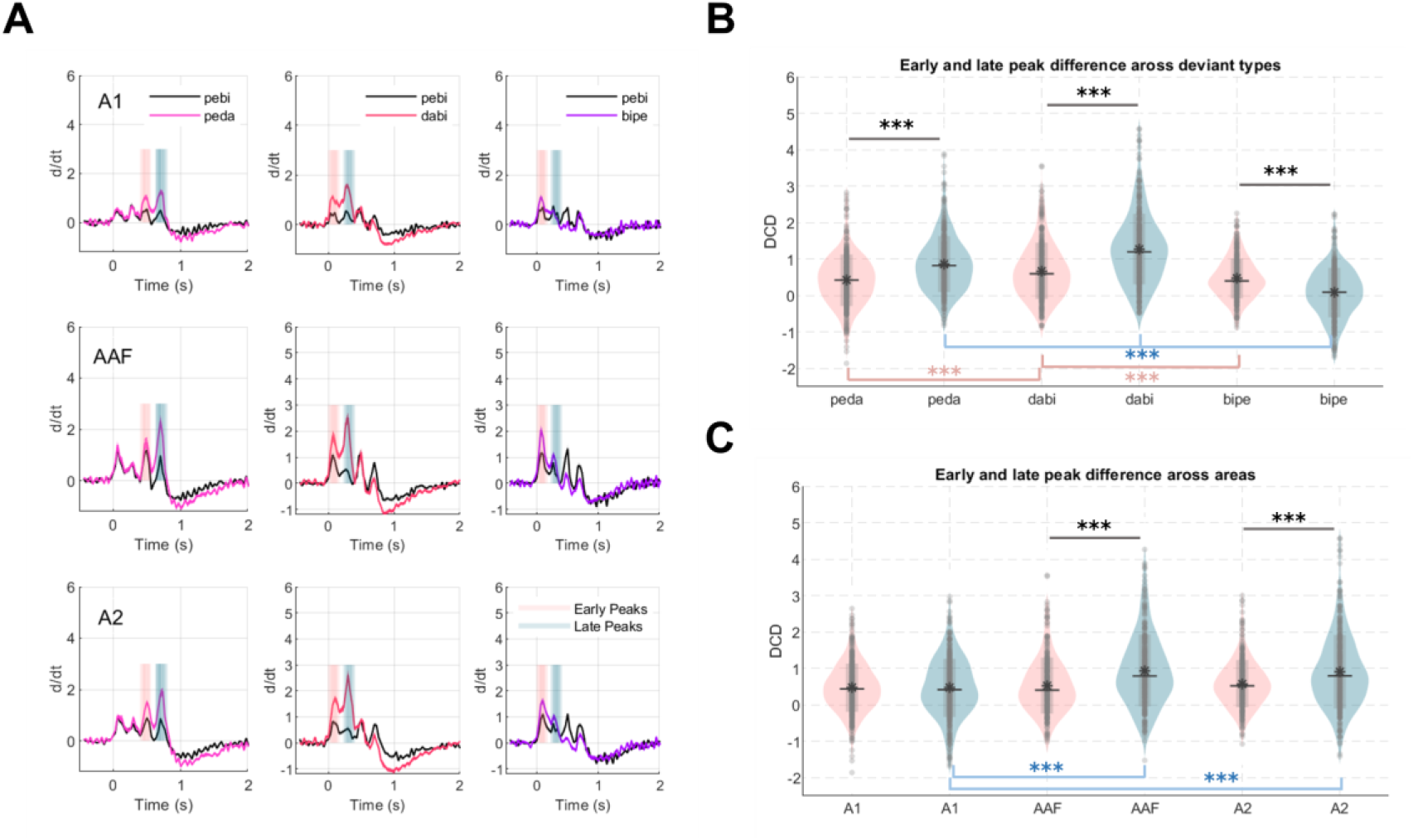
Temporal derivatives of calcium traces across deviant types and cortical areas. (A) The overall average of the first derivatives of the traces over A1, AAF and A2, plotted in the oddball condition and control condition, the pink bars and the blue bars indicate where we calculated the difference between the derivative values of deviants and standards for Fig. 5B&C. (B) Comparison between the difference of peaks at early and late time window for each of the different deviant types (peda, dabi, and bipe). Pink violin plots indicate the distribution of differences in early peak amplitude between deviants and standards. Blue violin plots indicate the corresponding late peaks differences. (C) Comparison between the difference of peaks at early and late time windows over different auditory areas (A1, AAF and A2). Pink indicates early peaks and blue indicates late peaks.

To test for peak differences between stimulus types in the time-resolved responses to deviants, we conducted a three-way ANOVA (cortical area * deviant type * early vs late DCD) on the DCD values calculated to the deviant elements in each pair as described above. These estimates were calculated separately for each cortical area. Based on these analyses, we found significant main effects of cortical area (*F* _(2,3086)_ = 13.5, p < 0.001), deviant type (*F* _(2,3086)_ = 570.38, p < 0.001), and “early vs late DCD” (*F* _(1,3086)_ = 150.38, p < 0.001).

Since previous studies had suggested that early and late parts of a response may contribute differently to deviant processing, we examined whether there were differences between the early and late DCD in different auditory cortical fields, and whether these depended on (i.e., interacted with) stimulus types. The results of our ANOVA suggests that this may indeed be the case, since we found a three-way interaction effect (cortical area * deviant type * early vs late DCD: *F* _(4,3086)_ = 7.95, p <0.001), as well as two-way interaction effects (deviant type * early vs late DCD: *F* _(2,3086)_ = 58.81, p < 0.001;cortical area * early vs late DCD: *F* _(2,2238)_ = 18.12, p < 0.001; cortical area * deviant type: *F* _(4,2238)_ = 6.42, p <0.001). Post-hoc tests revealed a differences between the early and late peak amplitudes for all deviant types (see Fig. 5B; ‘pe da’: Z = -7.845, p < 0.001; ‘da bi’: Z = -8.67, p < 0.001; ‘bi pe’: Z = 7.919, p < 0.001; all p values FDR corrected).

When comparing the early DCD across deviant types, we found that the early DCD was stronger for the substitution deviant ‘da bi’, in which the first syllable contains a deviant, than for the other substitution deviant ‘pe da’, in which the deviant only occurs during the second syllable, (rank-sum test, Z = 4.4, p < 0.001, FDR corrected). The early DCD for the substitution deviant ‘da bi’ was also stronger than for the transposition deviant ‘bi pe’ (Z = 3.834, p < 0.001, FDR corrected). No difference in early DCD was observed between deviant types ‘pe da’ and ‘bi pe’ (Z = -0.889, p = 0.374). For the late DCD, we also found that the deviant ‘da bi’ evoked stronger activity than ‘bi pe’ (Z = 16.9, p < 0.001, FDR corrected) and ‘pe da’ (Z = 6.185, p < 0.001, FDR corrected). However, here we also found that late DCD differed between the deviants ‘bi pe’ and ‘pe da’ (Z = 13.505, p < 0.001, FDR corrected). To summarize these results, there was no difference between the substitution deviant ‘pe da’ (the second syllable being a deviant) and the transposition deviant ‘bi pe’ in the early time window, but there was a substantial difference between these types of deviants in the late time window. Additionally, the DCD of the substitution deviant ‘da bi’ (the first syllable being a deviant) was larger than the other two deviants in both early and late time windows. These findings suggest that while the early DCD may be dependent on the preceding stimuli (‘da bi’ > ‘pe da’) as well as the transitional probability (‘da bi’ > ‘bi pe’), the late DCD may encode the content and novelty of the auditory stimuli (‘pe da’ > ‘bi pe’).

Turning to differences between the early vs. late DCD for each subdivision of the auditory cortex, post-hoc comparisons (rank-sum test, see Fig. 5C) showed that there were significant differences between the early vs. late DCD in A2 (Z = -4.389, p < 0.001, FDR corrected) and AAF (Z = -5.668, p < 0.001,FDR corrected), but not in A1 (Z = -0.112, p = 0.911). Furthermore, when analyzing the early DCD, no pairwise differences between A1, AAF, and A2 were found to be significant (all p > 0.05, FDR corrected). However, for the late DCD, the response amplitude was significantly smaller for A1 relative to AAF (Z = -5.882, p < 0.001, FDR corrected) and A2 (Z = - 5.461, p < 0.001, FDR corrected), with no significant difference between AAF and A2 (Z = 0.56, p = 0.576). These results suggest that differences in DCD between cortical areas are only observed for late calcium transients, and that late activity is most pronounced in higher-order regions.

### No mismatch response to the omitted sequence element across auditory fields

We observed no significant mismatch responses to the omission deviant in this study. The AUC of fluorescence to the omission deviant stimuli was found to be smaller than the standards in all auditory cortical fields. Specifically, when comparing the deviant with the pre-odd standard in A1, we found the AUC of fluorescence to the pre-odd standards is significantly larger than this to the omission pair ‘pe _’ (paired t-test, A1: t_41_ = 3.492, p = 0.001, AAF: t_25_ = 5.919, p < 0.001, A2: t_29_ = 9.807, p < 0.001, FDR corrected, see Fig. 6A). This may be explainable simply by the fact that the omission deviant comprised more silence and less acoustic energy than the standard, and a reduced cortical activation is therefore to be expected. We found similar results in all auditory fields when comparing the deviant with the first standard (A1: paired t-test, t_41_ = 10.845, AAF: t_25_=9.844, A2: t_29_=10.175, all p <0.001, FDR corrected, see Fig. 6A). In addition, when looking at the temporal derivatives of the calcium response, we observed no derivative peaks during the period of the omitted syllable in any of the all auditory fields (i.e. in the response to ‘pe _, we see transients only in response to the ‘pe’, not the ‘_’, see Fig. 6B). Thus, omissions of the second element of a standard pair did not seem to elicit robust omission responses that could be measured in the auditory cortex using calcium imaging.

**Figure 6.**
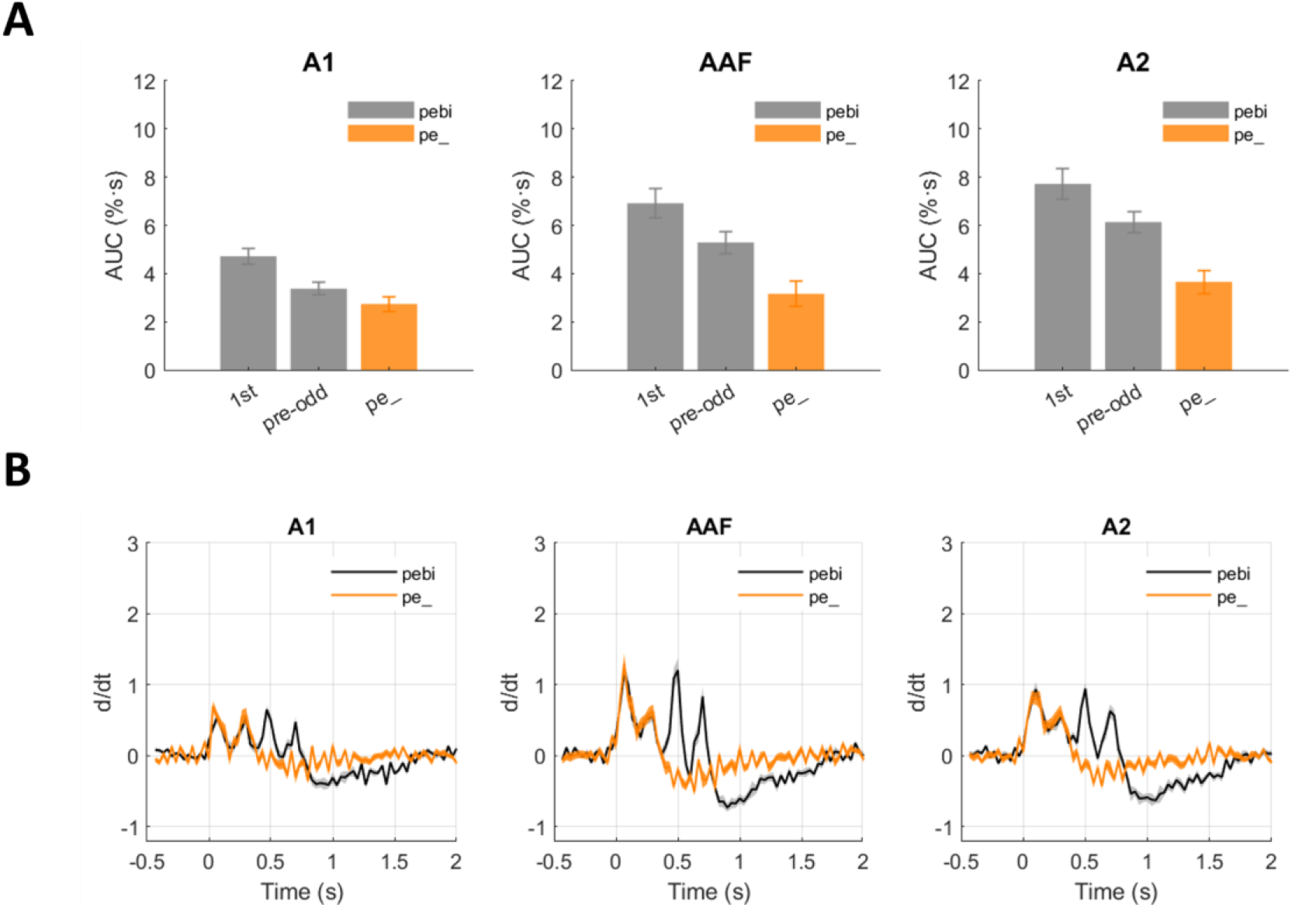
No mismatch responses found to complex stimulus omissions. (A) Comparison of calcium response to the first standard stimuli, pre-odd standard stimuli, and omission deviants across distinct auditory fields in the oddball condition. (B) The first derivatives of traces (dark: standard, yellow: pe_) from A1, AAF and A2 plotted for the oddball condition.

## Discussion

In this study, we examined whether the auditory cortical activity in awake mice shows sensitivity to violations of sequences based on simple deviance (single stimulus substitution) only, or whether it is also sensitive to more complex violations based on stimulus order (transposition). We found that widefield Ca++ responses in mouse cortex could encode both simple and global stimulus patterns, eliciting mismatch signals to the substitution deviants (‘pe bi’→ ‘pe da’ or ‘da bi’) and transposition (‘pe bi’→‘bi pe’) deviants (Fig. 4A), but not to omission deviants (‘pe bi’→‘pe _’; Fig. 6). For the substitution deviants, mismatch responses were more prominent during later parts of the response (Fig. 5B). These late mismatch responses were also larger in A2 than in A1 (Fig. 5C). Our results are thus consistent with previously articulated hypotheses regarding anatomical hierarchies of deviant processing (Parras et al., 2017), and functional distinctions between early and late calcium transients (Chen et al., 2015). Our results thus indicate that these ideas apply not just in the context of deviant processing for simple tones, but also to more complex, “chunked” stimuli such as bisyllabic sounds.

Mismatch responses likely comprise two partially dissociable phenomena: prediction error to the unexpected stimulus, and release from neural adaptation to the previously repeating standards. Consistent with previous studies in humans (Todorovic et al., 2011) and animals (Chen et al., 2015; Parras et al., 2017), we found a robust attenuation of responses to repetitively presented standards in all auditory fields (Fig. 3A). Similar effects have also been found in other modalities (Rao and Ballard, 1999; Shipp et al., 2013; Hamm and Yuste, 2016; Homann et al., 2017; Hamm et al., 2021). Based on the current experimental design, we cannot exclude the possibility that the gradual response attenuation found in our study was driven at least in part by passive adaptation (May and Tiitinen, 2010), since the temporal interval between stimulus pairs (∼2350 ms) was within the recovery range (about <10s) of synaptic depression (Ulanovsky et al., 2004). However, an increasing number of studies (Todorovic and de Lange, 2012; Wacongne et al., 2012; Auksztulewicz and Friston, 2016; Tang et al., 2018) suggest that response suppression to stimulus repetition is not only due to passive synaptic or spike rate adaptation, but it may at least in part also reflect a gradual reduction of a prediction error signal.

Mismatch responses to single deviant stimuli have been shown to follow a hierarchical gradient, such that stronger mismatch responses are observed in higher-order regions (Parras et al., 2017). Consistent with these studies, our analysis of responses to both substitution deviants (‘pe da’ and ‘da bi’) resulted in robust mismatch responses (Fig. 4A) which were larger in A2 than A1, suggesting a hierarchical gradient of deviance processing. Notably, a similar difference between auditory cortical regions (A2 > A1) was also identified when comparing the neural response (AUC) to ‘da bi’ deviants (in which the first element of the pair was surprising) with the first standards in the sequence (Fig. 4C), i.e., controlling for adaptation effects. However, no such effect was found for ‘pe da’ deviants (in which the second element of the pair was surprising, see Fig. 4B). This indicates that the serial position of the deviant element in the pair can modulate the prediction error, as the transitional probability between the surprising stimulus and the preceding stimulus is similar in both cases (‘pe bi’ followed by ‘da bi’, 3.93%; ‘pe bi’ followed by ‘pe da’, 3.66%). However, an alternative explanation might be that the neural habituation level is different in these two conditions due to differences in time intervals between the deviant stimulus and the preceding stimulus (‘pe da’: 145 ms gap between the expected ‘pe’ and the surprising ‘da’; ‘da bi’: 2350 ms gap between the pre-odd standard ‘bi’ and the surprising ‘da’). While in this study we introduced differences in time intervals to facilitate chunking, future studies should investigate similar serial effects in isochronous sequences.

Crucially, our results extend the notion of a hierarchical gradient of mismatch responses also to more complex prediction violations, based on stimulus order. Specifically, both A2 and AAF (but not A1) elicited MMRs to the transposition deviants when compared against the immediately preceding standards. However, in this case we also identified a major contribution of repetition suppression to the mismatch signal, as no difference was found when comparing the deviants against the first standard. Furthermore, the MMRs were found to be much weaker for the transposition deviants than for substitution deviants, although the transition probability between the surprising syllable and the preceding standard syllable was matched between deviant types (‘pe bi’→‘bi’, 3.93%, ‘pe bi’→‘da’, 3.93%). Therefore, while we found evidence for neural sensitivity to stimulus order violations in higher-order cortical regions, unexpected substitutions constitute a more influential cue for deviant processing in this study. It should also be noted that a recent electrophysiology study (Parras et al., 2021) suggested that the PAF (posterior auditory field) produces an even more robust prediction error response to single deviants than A2. Future studies on responses to complex substitution deviants should expand the imaging range of cortical regions to include the PAF, to test whether the gradient of mismatch responses extends further beyond A2, and whether PAF is more sensitive to transposition deviants.

While calcium traces have relatively slow dynamics, analyzing their temporal derivative can yield dissociable early and late peaks, which have previously been mapped onto adaptation and deviance detection respectively (Chen et al., 2015). However, it is not clear whether the early and late peaks can also be differentially modulated by deviance processing across the cortical hierarchy, although a previous electrophysiology study showed that higher-order regions have relatively longer mismatch response latencies (Nieto-Diego and Malmierca, 2016). Our analysis revealed that late peak amplitudes were greater in A2/AAF than in A1, while there was no difference between cortical areas at the earlier time window. Furthermore, we observed an enhancement of late responses (relative to early responses) in A2/AAF, but no such difference in A1. This suggests that higher-order regions may signal mismatch responses at relatively longer latencies than hierarchically lower areas. However, based on the timing of our stimuli, we cannot exclude the possibility that offset responses may also contribute to the late response, which may also explain the differences between AAF and A1, as AAF might exhibit stronger off responses than A1 (Solyga and Barkat, 2019, 2021)(Liu et al., 2019). Turning to the functional distinction between early and late calcium transients, we found that the late deviant response was stronger than the early deviant response for both substitution deviants (Fig. 5B), suggesting that the late response contributed more to the overall mismatch response (prediction error signal) in both deviants than the early response. By contrast, the early response was stronger than the late response for the transposition deviants, but weaker than the early response for those substitution deviants where the first pair element was unexpected. This suggests that the early response may reflect release from adaptation to the preceding repeated pair, and the late response may reflect novelty of the deviant stimulus content.

In contrast, responses to omission deviants (Fig. 6A) showed no robust mismatch response to the omitted stimulus. While some previous studies did identify calcium ‘echo’ signals following isochronous sequence termination (Li et al., 2017; Wang et al., 2018), they necessitated a large number of standard repetitions and were facilitated by previous behavioral training. Our null findings suggest a number of non-exclusive possibilities. First, it could be that omission responses are not robustly encoded in the widefield calcium activity (analyzed here) or spiking activity mainly from excitatory neurons in the superficial layers (to which wide-field imaging is most sensitive) (Waters, 2020), but rather in population activity (Auksztulewicz et al., 2022) potentially reflecting dendritic currents (Guerguiev et al., 2017), or in other types of neurons. This possibility is consistent with a recent calcium imaging study in the visual cortex, where neural activity following omissions was observed in L2/3 inhibitory neurons, but not in L2/3 excitatory neurons (Garrett et al., 2020). Our temporal derivative results (see Fig. 6B) indeed suggested that there were no calcium events during the omitted stimulus (the second element of a pair), while calcium events for pair-final elements were found in other types of stimuli and were modulated by the deviant type (see Fig. 5A). Another possibility pertains to stimulus characteristics, as recent evidence from a primate study (Jiang et al., 2022) showed that the omission response was more pronounced when the omitted tone was presented in a rhythmic condition at a fixed rate. In contrast, the omission in our study was not at a fixed rate, and pertained to a missing element of a pair, rather than an entire missing standard as in previous studies (Li et al., 2017; Wang et al., 2018). Taken together, more evidence is needed to elucidate the types of neurons and population mechanisms underlying omission signaling.

In summary, we show that mice can encode the deviants based on single stimulus substitutions, but also based on order transpositions. Higher-order auditory regions (A2) showed larger MMRs than primary regions (A1), especially at the late time window, which extends previous findings in traditional oddball paradigms based on single deviants, showing an increase of mismatch signaling along the auditory processing pathway (Parras et al., 2017). Our findings thus show that the mouse auditory cortex is a suitable model also for deviant processing based on more complex transitional probabilities.

## Acknowledgments

Supported by NIH R01DC009607 (POK), NIH RO1DC017785 (POK), Hong Kong General Research Fund 11100518 and an EU/HK Research Grants Council grant (JS).

